# APC/C^FZR-1^ Controls SAS-5 Levels to Regulate Centrosome Duplication in *Caenorhabditis elegans*

**DOI:** 10.1101/186148

**Authors:** Jeffrey C. Medley, Lauren E. DeMeyer, Megan M. Kabara, Mi Hye Song

## Abstract

As the primary microtubule-organizing center, centrosomes play a key role in establishing mitotic bipolar spindles that secure correct transmission of genomic content. For the fidelity of cell division, centrosome number must be strictly controlled by duplicating only once per cell cycle. Proper levels of centrosome proteins are shown to be critical for normal centrosome number and function. Overexpressing core centrosome factors leads to extra centrosomes, while depleting these factors results in centrosome duplication failure. In this regard, protein turnover by the ubiquitin-proteasome system provides a vital mechanism for the regulation of centrosome protein levels. Here, we report that FZR-1, the *Caenorhabditis elegans* homolog of Cdh1/Hct1/Fzr, a co-activator of the anaphase promoting complex/cyclosome (APC/C), an E3 ubiquitin ligase, functions as a negative regulator of centrosome duplication in the *Caenorhabditis elegans* embryo. During mitotic cell division in the early embryo, FZR-1 is associated with centrosomes and enriched at nuclei. Loss of *fzr-1* function restores centrosome duplication and embryonic viability to the hypomorphic *zyg-1(it25)* mutant, in part, through elevated levels of SAS-5 at centrosomes. Our data suggest that the APC/C^FZR-1^ regulates SAS-5 levels by directly recognizing the conserved KEN-box motif, contributing to proper centrosome duplication. Together, our work shows that FZR-1 plays a conserved role in regulating centrosome duplication in *Caenorhabditis elegans*.

## INTRODUCTION

The centrosome is a small, non-membranous organelle that serves as the primary microtubule-organizing center in animal cells. Each centrosome consists of a pair of barrel-shaped centrioles that are surrounded by a network of proteins called pericentriolar material (PCM). During mitosis, two centrosomes organize bipolar spindles that segregate genomic content equally into two daughter cells. Thus, tight control of centrosome number is vital for the maintenance of genomic integrity during cell division, by restricting centrosome duplication to once and only once per cell cycle. Erroneous centrosome duplication results in aberrant centrosome number that leads to chromosome missegregation and abnormal cell proliferation and is associated with human disorders including cancers and microcephaly (Nigg and Stearns 2011; Gönczy 2015).

In the nematode *C. elegans*, extensive studies identified a set of core centrosome factors that are absolutely essential for centrosome duplication: the protein kinase ZYG-1 and the coiled-coil proteins SPD2, SAS-4, SAS-5 and SAS-6 (O’Connell *et al.* 2001; Kirkham *et al.* 2003; Leidel and Gönczy 2003; Dammermann *et al.* 2004; Delattre *et al.* 2004; Kemp *et al.* 2004; Pelletier *et al.* 2004; Leidel *et al.* 2005). SPD-2 and ZYG-1 localize early to the site of centriole formation and are required for the recruitment of the SAS-5/SAS-6 complex that sequentially recruits SAS-4 to the centriole (Delattre *et al.* 2006; Pelletier *et al.* 2006). These key factors are also present in other animal systems, suggesting highly conserved evolutionary mechanisms in centrosome duplication. For instance, the human genome contains homologs of the five centrosome factors found in *C. elegans*, Cep192/SPD-2 (Zhu *et al.* 2008), Plk4/ZYG-1 (Habedanck *et al.* 2005), STIL/SAS-5 (Arquint *et al.* 2012), HsSAS-6/SAS-6 (Leidel *et al.* 2005) and CPAP/SAS-4 (Kleylein-Sohn *et al.* 2007; Tang *et al.* 2009) and all these factors are shown to play a critical role in centrosome biogenesis (Fu *et al.* 2015; Gönczy 2015).

Maintaining the proper levels of centrosome proteins is critical for normal centrosome number and function (Kleylein-Sohn *et al.* 2007; Strnad *et al.* 2007; Rogers *et al.* 2009; Tang *et al.* 2009; Holland *et al.* 2010; Brownlee *et al.* 2011; Puklowski *et al.* 2011; Song *et al.* 2011; Tang *et al.* 2011; Meghini *et al.* 2016; Levine *et al.* 2017). In light of this, protein turnover by proteolysis provides a key mechanism for regulating the abundance of centrosome factors. A mechanism regulating protein levels is their degradation by the 26S proteasome that catalyzes the proteolysis of poly-ubiquitinated substrates (Livneh *et al.* 2016). The anaphase promoting complex/cyclosome (APC/C) is a multi-subunit E3 ubiquitin ligase that targets substrates for degradation (Acquaviva and Pines 2006; Peters 2006; Chang and Barford 2014). The substrate specificity of the APC/C is directed through the sequential, cell cycle-dependent activity of two co-activators, Cdc20/Fzy/FZY-1 (Hartwell and Smith 1985; Dawson *et al.* 1995; Kitagawa *et al.* 2002) and Cdh1/Fzr/Hct1/FZR-1 (Schwab *et al.* 1997; Sigrist and Lehner 1997; Visintin *et al.* 1997; Fay *et al.* 2002). During early mitosis Cdc20 acts as co-activator of the APC/C, and Cdh1 functions as co-activator to modulate the APC/C-dependent events at late mitosis and in G1 (Irniger and Nasmyth 1997; Visintin *et al.* 1997; Fang *et al.* 1998; Prinz *et al.* 1998; Shirayama *et al.* 1998). Upregulated targets in Cdh1-deficient cells are shown to be associated with the genomic instability signature of human cancers and show a high correlation with poor prognosis (Carter *et al.* 2006; Garcia-Higuera *et al.* 2008). Furthermore, a mutation in SIL/STIL (a human homolog of SAS-5) linked to primary microcephaly (MCPH; Kumar *et al.* 2009) results in deletion of the Cdh1-dependent destruction motif (KEN-box), leading to deregulated accumulation of STIL protein and centrosome amplification (Arquint and Nigg 2014). In *Drosophila*, the APC/C^Fzr/Cdh1^ directly interacts with Spd2 through KEN-box recognition and targets Spd2 for degradation (Meghini *et al.* 2016). Therefore, the APC/C^Cdh1/Fzr/Hct1^ plays a critical role in regulating the levels of key centrosome duplication factors in mammalian cells and flies.

In *C. elegans*, FZR-1 has been shown to be required for fertility, cell cycle progression and cell proliferation during embryonic and postembryonic development via synthetic interaction with *lin-35/Rb* (Fay *et al.* 2002; The *et al.* 2015). However, the role of FZR-1 in centrosome assembly has not been described. In this study, we molecularly identified *fzr-1* as a genetic suppressor of *zyg-1.* Our results suggest that APC/C^FZR-1^ negatively regulates centrosome duplication, in part, through proteasomal degradation of SAS-5 in a KEN-box dependent fashion. Therefore, FZR-1, the *C. elegans* homolog of Cdh1/Hct1/Fzr, plays a conserved role in centrosome duplication.

## MATERIALS AND METHODS

### *C. elegans* strains and genetics

A full list of *C. elegans* strains used in this study is listed in Table S1. All strains were derived from the wild-type Bristol N2 strain using standard genetic methods (Brenner 1974; Church *et al.* 1995). Strains were maintained on MYOB plates seeded with *E. coli* OP50 and grown at 19° unless otherwise indicated. The *fzr-1∷gfp∷3xflag* construct containing 21.6Kbp of the *fzr-1* 5’UTR and 6Kbp of the *fzr-1* 3’UTR was acquired from TransgenOme (construct number: 7127141463160758 F11, Sarov *et al.* 2012), which was used to generate the transgenic line, MTU10, expressing C-terminal GFP-tagged FZR-1. For the generation of N-terminal GFP-tagged FZR-1 (OC190), we used Gateway cloning (Invitrogen, Carlsbad, CA, USA) to generate the construct. Coding sequence of *fzr-1* was PCR amplified from the cDNA clone yk1338f2, and cloned into pDONR221 (Invitrogen) and then the resulting pDONR construct was recombined into pID3.01 (pMS9.3), which is driven by the *pie-1* promotor. The transgenes were introduced into worms by standard particle bombardment (Praitis *et al.* 2001). For embryonic viability and brood size assays, individual L4 animals were transferred to clean plates and allowed to self-fertilize for 24 hours at the temperatures indicated. For brood size assays, this was repeated until animals no longer produced fertilized embryos. Progeny were allowed at least 24 hours to complete embryogenesis before counting the number of progeny. The *fzr-1(RNAi)* experiments were performed by RNAi soaking (Song *et al.* 2008). To produce dsRNA for RNAi soaking, we amplified a DNA template from the cDNA clone yk1338f2 using the primers 5’-ATGGATGAGCAACCGCC-3’ and 5’-GCACTGTACGTAAAAAGTGATC-3’ that contained a T7 promoter sequence at their 5’ ends. *In vitro* transcription was performed using the T7-MEGAscript kit (Thermo-Fisher, Hanover park, IL, USA). L4 animals were soaked overnight in M9 buffer containing either 0.1-0.4 mg dsRNA/mL or no dsRNA (control).

### Mapping and molecular identification of *szy-14*

We mated *zyg-1(it25) dpy-10(e128) szy-14(bs31) unc-4(e120)* hermaphrodites with Hawaiian, CB4856 males for single-nucleotide polymorphism mapping (Song *et al.* 2008), and isolated a total of 104 independent Dpy-nonUnc from the F2 generation. After establishing homozygous recombinant lines, we scored for the presence of *szy-14(bs31)* based on the suppression of the *zyg-1(it25)* mutant (additionally, reduced brood size; (Fay *et al.* 2002). In parallel, we used *zyg-1(it25), zyg-1(it25) dpy-10(e128), zyg-1(it25) dpy-10(e128) szy-14(bs31) unc-4(e120), zyg-1(it25) szy-14(bs31)*, and *zyg-1(it25) szy-14(bs31) unc-4(e120)* as controls. For the molecular identification of the mutation, we sequenced several candidate genes (*nos-3, kin-15, kin-16, wee-1.1, wee-1.3*, and *fzr-1*) located within an interval on chromosome II. For sequencing the *fzr-1* gene, we used the following primers: Forward 5’-TCTTGTTTCTGGTGGAGGT-3’ and Reverse 5’- ACACGATACTGATGCCCAA-3’ for the *bs31* suppressor, and Forward 5’- ATGGATGAGCAGCAACCGCC-3’ and Reverse 5’-CAAGCTTGAGCTGTTGG-3’ for the *bs38* suppressor. Purified PCR amplicons were sequenced and aligned to the ORF, ZK1307.6 to identify the nucleotide substitution.

### CRISPR/CAS-9 mediated genome editing

For genome editing, we used the co-CRISPR technique as previously described in *C. elegans* (Arribere *et al.* 2014; Paix *et al.* 2015). In brief, we microinjected N2 and *zyg-1(it25)* animals using a mixture containing recombinant SpCas9 (Paix *et al.* 2015), crRNAs targeting *sas-5* and *dpy-10* at 0.4-0.8μg/μL, tracrRNA at 12μg/μL, and single-stranded DNA oligonucleotides to repair *sas-5* and *dpy-10* at 25-100ng/μL. Microinjection was performed using the XenoWorks microinjector (Sutter Instruments, Novato, CA, USA) with a continuous pulse setting at 400-800 hPa. All RNA and DNA oligonucleotides used in this study were synthesized by Integrated DNA Technologies (IDT, Coralville, IA, USA) and are listed in Table S2. As we were unable to engineer a silent mutation into the PAM sequence used by the *sas-5* crRNA, we introduced six silent mutations to *sas-5* (aa 201-206; Figure 5A) by mutating 8 out of 20 the nucleotides that comprise the *sas-5* crRNA, in order to disrupt Cas9 recognition after homology-directed repair. After injection, animals were allowed to produce F1 progeny that were monitored for the presence of *dpy-10(cn64)/+* rollers. To identify the *sas-5^KEN-to-3A^* mutation, we extracted genomic DNA from broods containing the highest frequency of F1 rollers. Using the primers, forward: 5’-TGCCCAAAATACGACAACG-3’ and reverse: 5’-TACACTACTCACGTCTGCT-3’, we amplified the region of *sas-5* containing the KEN-box sequence. As the repair template for the *sas-5^KEN-to-3A^* mutation introduces an *Hpy8*I restriction enzyme (NEB, Ipswich, MA, USA) cutting site, we used an *Hpy8*I enzyme digestion to test for the introduction of our targeted mutation. After isolating homozygotes based on the *Hpy8*I cutting, we confirmed the *SAS-5^KEN-to-3A^* mutation by genomic DNA sequencing. Sequencing revealed that several lines were homozygous for the *SAS-5^KEN-to-3A^* mutation (Table S1, Figure 5A). However, the strain MTU14, contained all of the silent mutations that we designed to disrupt Cas9 recognition without affecting the KEN-box (Table S1, Figure 5A). Thus, we used MTU14 as a control for our assays.

### Cytological analysis

To perform immunostaining, the following antibodies were used at 1:2,000-3,000 dilutions: α-Tubulin (DM1a; Sigma, St-Louis, MO, USA), α-GFP: IgG_1_K (Roche, Indianapolis, IN, USA), α-ZYG-1 (Stubenvoll *et al.* 2016), α-TBG-1(Stubenvoll *et al.* 2016), α-SAS-4 (Song *et al.* 2008), α-SAS-5 (Medley *et al.* 2017), and Alexa Fluor 488 and 561 (Invitrogen, Carlsbad, CA, USA) as secondary antibodies. Confocal microscopy was performed as described (Stubenvoll *et al.* 2016) using a Nikon Eclipse Ti-U microscope equipped with a Plan Apo 60 x 1.4 NA lens, a Spinning Disk Confocal (CSU X1) and a Photometrics Evolve 512 camera. Images were acquired using MetaMorph software (Molecular Devices, Sunnyvale, CA, USA). MetaMorph was used to draw and quantify regions of fluorescence intensity and Adobe Photoshop CS6 was used for image processing. To quantify centrosomal SAS-5 signals, the average intensity within an 8-pixel (1 pixel = 0.151 μm) diameter region was measured within an area centered on the centrosome and the focal plane with the highest average intensity was recorded. Centrosomal TBG-1 (γ-tubulin) levels were quantified in the same manner, except that a 25-pixel diameter region was used. For both SAS-5 and TBG-1 quantification, the average fluorescence intensity within a 25-pixel diameter region drawn outside of the embryo was used for background subtraction.

### Immunoprecipitation (IP)

Embryos were collected from gravid worms using hypochlorite treatment (1:2:1 ratio of M9 buffer, 5.25% sodium hypochlorite and 5M NaCl), washed with M9 buffer five times and frozen in liquid nitrogen. Embryos were stored at -80° until use. IP experiment using α-GFP were performed following the protocol described previously (Stubenvoll *et al.* 2016). 20 μL of Mouse-α-GFP magnetic beads (MBL, Naka-ku, Nagoya, Japan) were used per reaction. The α-GFP beads were prepared by washing twice for 15 minutes in PBST (PBS; 0.1% Triton-X), followed by a third wash in 1x lysis buffer (50 mM HEPES [pH 7.4], 1mM EDTA, 1mM MgCl__2__, 200 mM KCl, and 10% glycerol (v/v)) (Cheeseman *et al.* 2004). Embryos were suspended in 1 x lysis buffer supplemented with complete protease inhibitor cocktail (Roche, Indianapolis, IN, USA) and MG132 (Tocris, Avonmouth, Bristol, UK). The embryos were then milled for three minutes at 30 Hz using a Retsch MM 400 mixer-mill (Verder Scientific, Newtown, PA, USA). Lysates were sonicated for three minutes in ice water using an ultrasonic bath (Thermo-Fisher, Hanover Park, IL, USA). Samples were spun at 45,000RPM for 45 minutes using a Sorvall RC M120EX ultracentrifuge (Thermo-Fisher, Hanover Park, IL, USA). The supernatant was transferred to clean microcentrifuge tubes. Protein concentration was quantified using a NanoDrop spectrophotometer (Thermo-Fisher, Hanover Park, IL, USA) and equivalent amount of total proteins was used for each reaction. Samples and α-GFP beads were incubated and rotated for one hour at 4°C and then washed five times for five minutes using PBST (PBS + 0.1% Triton-X 100). Samples were resuspended in 20 μL of a solution containing 2X Laemmli Sample Buffer (Sigma, St-Louis, MO, USA) and 10% β-mercaptoethanol (v/v), then boiled for five minutes. For protein input, 5 μL of embryonic lysates were diluted using 15 μL of a solution containing 2X Laemmli Sample Buffer and 10% β-mercaptoethanol (v/v) and boiled for 5 minutes before fractionating on a 4-12% NuPAGE Bis-Tris gel (Invitrogen, Carlsbad, CA, USA).

### Western Blotting

For western blotting, samples were sonicated for five minutes and boiled in a solution of 2X Laemmli Sample Buffer and 10% β-mercaptoethanol before being fractionated on a 4-12% NuPAGE Bis-Tris gel (Invitrogen, Carlsbad, CA, USA). The iBlot Gel Transfer system (Invitrogen, Carlsbad, CA, USA) was then used to transfer samples to a nitrocellulose membrane. The following antibodies were used at 1:3,000-10,000 dilutions: α-Tubulin: α-Tubulin (DM1a; Sigma, St-Louis, MO, USA), α-GFP: IgG_1_K (Roche, Indianapolis, IN, USA), α-SAS-5 (Song *et al.* 2011) and α-TBG-1 (Stubenvoll *et al.* 2016). IRDye secondary antibodies (LI-COR Biosciences, Lincoln, NE, USA) were used at a 1:10,000 dilution. Blots were imaged using the Odyssey infrared scanner (LI-COR Biosciences, Lincoln, NE, USA), and analyzed using Image Studio software (LI-COR Biosciences, Lincoln, NE, USA).

### Statistical Analysis

All *p*-values were calculated using two-tailed t-tests assuming equal variance among sample groups. Statistics are presented as Average ± standard deviation (SD) unless otherwise specified. Data were independently replicated at least three times for all experiments and subsequently analyzed for statistical significance.

### Data Availability

All strains used in this study are available upon request. The following supplemental materials are uploaded.

Figure S1. Centrosome-associated TBG-1 levels are unaffected in *fzr-1(bs31)* and *sas-5^KEN-to-3A^* mutant embryos.

Figure S2. Brood size in *sas-5*^*KEN-to-3A*^ and *fzr-1(bs31)* mutants.

Figure S3. SAS-5 levels are increased in *sas-5*^*KEN-to-3A*^ mutants.

Table S1. List of strains used in this study.

Table S2. List of oligonucleotides used for CRISPR/Cas9 genome editing.

## RESULTS AND DISCUSSION

### The *szy-14* mutation restores centrosome duplication to *zyg-1(it25)* mutants

Through a genetic suppressor screen (Kemp *et al.* 2007), the *szy-14* (<u>s</u>uppressor of *<u>zy</u>g-1*) gene was originally identified that restores embryonic viability of the partial loss-of-function *zyg-1(it25)* mutant. The *zyg-1(it25)* mutant embryo grown at the restrictive temperature (24°) fails to duplicate centrosomes during the first cell cycle, resulting in monopolar spindles at the second mitosis and 100% embryonic lethality (O’Connell *et al.* 2001). A complementation test identified two alleles, *szy-14(bs31)* and *szy-14(bs38)*, of the *szy-14* mutation that partially restore the embryonic viability of *zyg-1(it25)* but show slow growth phenotype without obvious cytological defects, indicating that the *szy-14* gene is not essential for embryonic viability (Table 1, (Kemp *et al.* 2007).

**Table 1.**
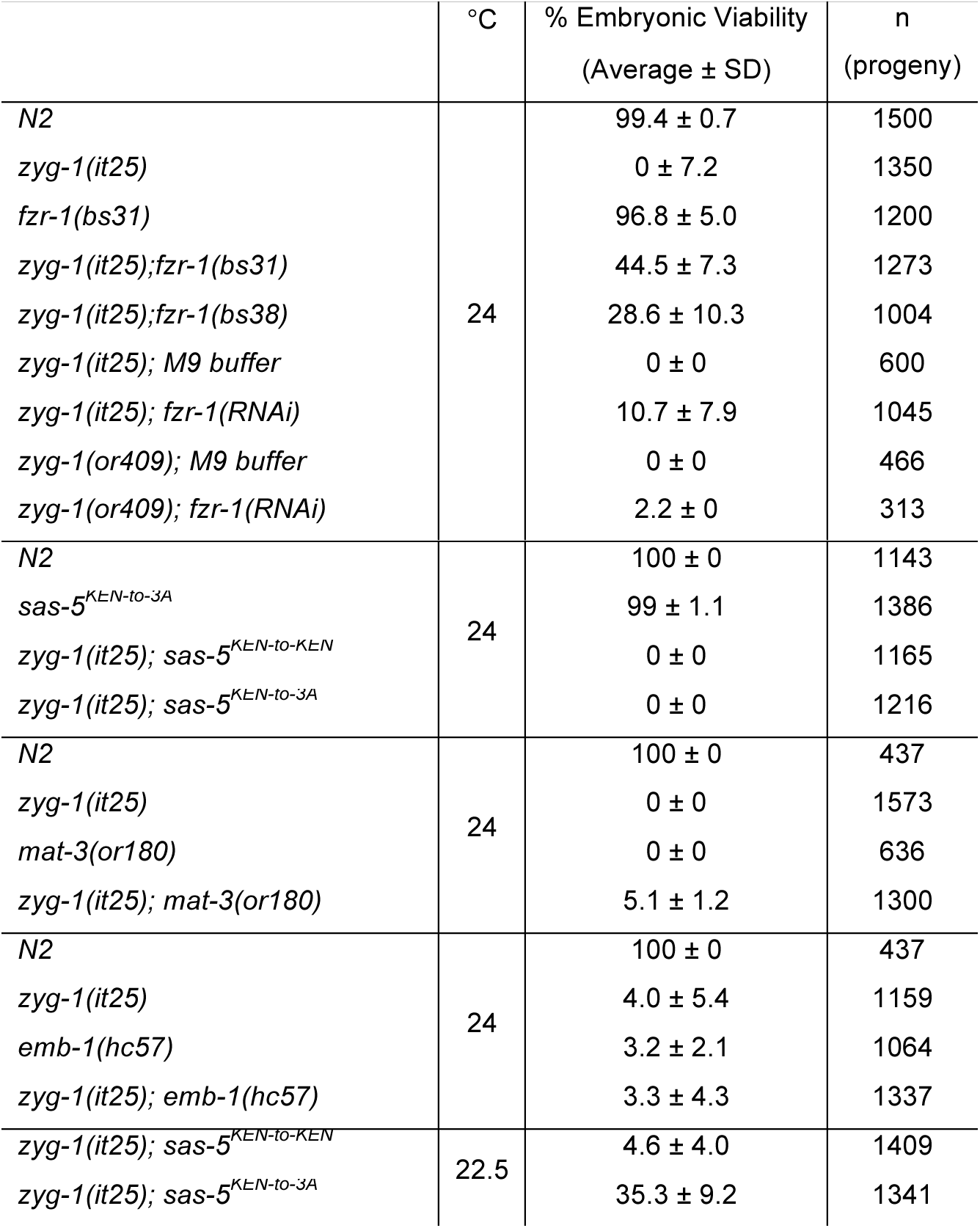
Genetic Analysis

Given that ZYG-1 is essential for proper centrosome duplication (O’Connell *et al.* 2001), we speculated that the *szy-14* mutation might suppress the embryonic lethality of *zyg-1(it25)* mutants via restoration of centrosome duplication. To examine centrosome duplication events, we quantified the percentage of bipolar spindles at the second mitosis, which indicates successful centrosome duplication during the first cell cycle (Figure 1A and B). At the restrictive temperature 24°, both double mutant embryos, *zyg-1(it25); szy-14(bs31)* (79.9±22.0%) and *zyg-1(it25); szy-14(bs38)* (51.4±24.4%) produced bipolar spindles at a significantly higher rate, compared to *zyg-1(it25)* single mutant embryos (3.3±4.4%) (Figure 1B). Our observation suggests that the *szy-14* mutation restores centrosome duplication in *zyg-1(it25)* embryos, thereby restoring embryonic viability to *zyg-1(it25)* mutants.

**Figure 1.**
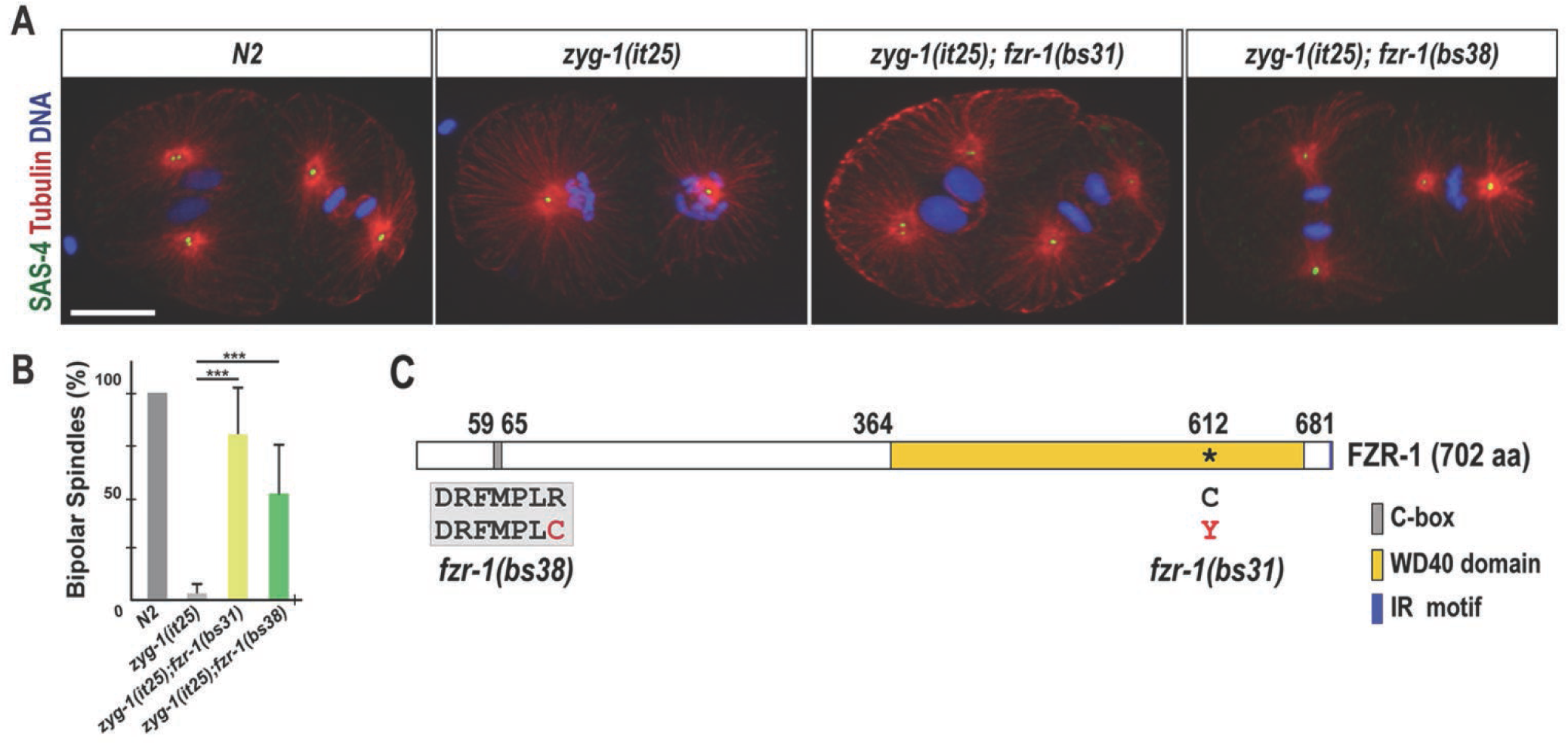
*fzr-1* mutations restore bipolar spindle formation to *zyg-1(it25)*. (A) Embryos grown at 24° stained for centrosomes (SAS-4), microtubules and DNA, illustrating mitotic spindles at the second mitosis. In *zyg-1(it25); fzr-1(bs31)* and *zyg-1(it25); fzr-1(bs38)* double mutant embryos, bipolar spindle formation is restored, whereas the *zyg-1(it25)* mutant embryo forms monopolar spindles. The *N2* embryo is shown as a wild-type control that shows bipolar spindles. Bar, 10 μm. (B) Quantification of bipolar spindle formation during the second cell cycle. At the restrictive temperature (24°), a great majority of *zyg-1(it25)* mutant embryos form monopolar spindles (3.3±4.4% bipolar spindles, n=660 blastomeres). In contrast, bipolar spindle formation is restored in *zyg-1(it25); fzr-1(bs31)* (79.9±22.0% bipolar spindles, n=276 blastomeres, *p*<0.001) and *zyg-1(it25); fzr-1(bs38)* (51.4±24.4% blastomeres, n=404 blastomeres, *p*<0.001) double mutants. Wild-type (*N2*) embryos invariably assemble bipolar spindles (100% bipolar spindles, n=600 blastomeres). Average values are presented. Error bars represent standard deviation (SD). \*\*\**p*<0.001 (two-tailed t-test). (C) Schematic of FZR-1 protein structure illustrates functional domains and the location of the missense mutations: R65C within the C-box in the *fzr-1(bs38)* mutant, and C612Y within WD40 domain in the *fzr-1(bs31)* mutant allele.

### Molecular identification of *szy-14*

The *szy-14* gene was initially mapped to the right arm of chromosome II between the morphological markers *dpy-10* and *unc-4* (Kemp *et al.* 2007). Using fine physical mapping, we located *szy-14* to an interval of 161-Kb (ChrII: 9621265..9782352; Wormbase.org) that contains several known cell cycle regulators. Based on the genetic map position of the *szy-14* suppressor, we sequenced candidate genes within this interval to detect any mutations in *szy-14* mutants. Sequencing revealed that *szy-14(bs38)* mutants contain a single substitution (C-to-T) in exon 2, and *szy-14(bs31)* mutants carry a mutation (G-to-A) in exon 5 of the ORF ZK1307.6 that corresponds to the *fzr-1* gene. Consistently, inhibiting FZR-1 by RNAi soaking partially restores embryonic viability in both *zyg-1(it25)* and *zyg-1(or409)* mutant alleles (Table 1), indicating that loss-of-function of *fzr-1* leads to the restoration of embryonic viability to the *zyg-1* mutants. Together, we determined that the *bs31* and *bs38* mutations are alleles of the *fzr-1* gene. Hereafter, we refer to *szy-14(bs31)* and *szy-14(bs38)* mutants as *fzr-1(bs31)* and *fzr-1(bs38)* mutants, respectively.

*fzr-1* encodes a conserved co-activator of the anaphase promoting complex/cyclosome (APC/C), the *C. elegans* homolog of Cdh1/Hct1/Fzr (Schwab *et al.* 1997; Sigrist and Lehner 1997; Visintin *et al.* 1997; Fay *et al.* 2002). The APC/C is an E3 ubiquitin ligase that orchestrates the sequential degradation of key cell cycle regulators during mitosis and early interphase (Song and Rape 2008). As part of this process, specific activators modulate the APC/C activity in different phases of mitosis. Specifically, FZR-1/Cdh1 modulates the APC/C at late mitosis and events in G1 during the time when centrosome duplication occurs. In each of the *fzr-1* mutant alleles, the single substitution leads to a missense mutation (Figure 1C). The *fzr-1(bs31)* mutation results in a missense mutation (C612Y) within the conserved WD40 repeat domain that is known to be involved in protein-protein interactions and is important for substrate recognition (Kraft *et al.* 2005; He *et al.* 2013). The *fzr-1(bs38)* mutation produces a missense mutation (R65C) at the conserved C-box of FZR-1. The C-box is known to be crucial for the physical interaction between FZR-1 and other APC/C subunits (Schwab *et al.* 2001; Thornton *et al.* 2006; Chang *et al.* 2015; Zhang *et al.* 2016). Thus, both *fzr-1(bs31)* and *fzr-1(bs38)* mutations appear to affect conserved domains that are critical for the function of the APC/C complex, suggesting that FZR-1 might regulate centrosome duplication through the APC/C complex.

### FZR-1 localizes to nuclei and centrosomes during early cell division

To determine where FZR-1 might function during the early cell cycle, we produced two independent transgenic strains that express FZR-1 tagged with GFP at the N-or C-terminus (see method and materials). To label microtubules, we mated GFP-tagged FZR-1 transgenic animals with the mCherry∷β-tubulin expressing line, and performed 4D time-lapse movies to observe subcellular localization of FZR-1 throughout the first cell cycle (Figure 2A). Confocal imaging illustrates that during interphase and early mitosis, FZR-1 is highly enriched at the nuclei. After the nuclear envelope breaks down (NEBD), FZR-1 diffuses to the cytoplasm and reappears to the nuclei at late mitosis when the nuclear envelop reforms. After NEBD, FZR-1 becomes apparent at spindle microtubules, and centrosomes that co-localize with SPD-2, a centrosome protein (Figure 2B). Both GFP-tagged FZR-1 transgenic embryos exhibit similar subcellular distributions, except a slight difference in fluorescent intensity (not shown). Our observations suggest that *C. elegans* FZR-1 might direct APC/C activity at centrosomes during late mitosis in early embryos, which is consistent with the role of FZR-1 as the co-activator of the APC/C at late mitosis in other organisms (Raff *et al.* 2002; Zhou *et al.* 2003; Meghini *et al.* 2016).

**Figure 2.**
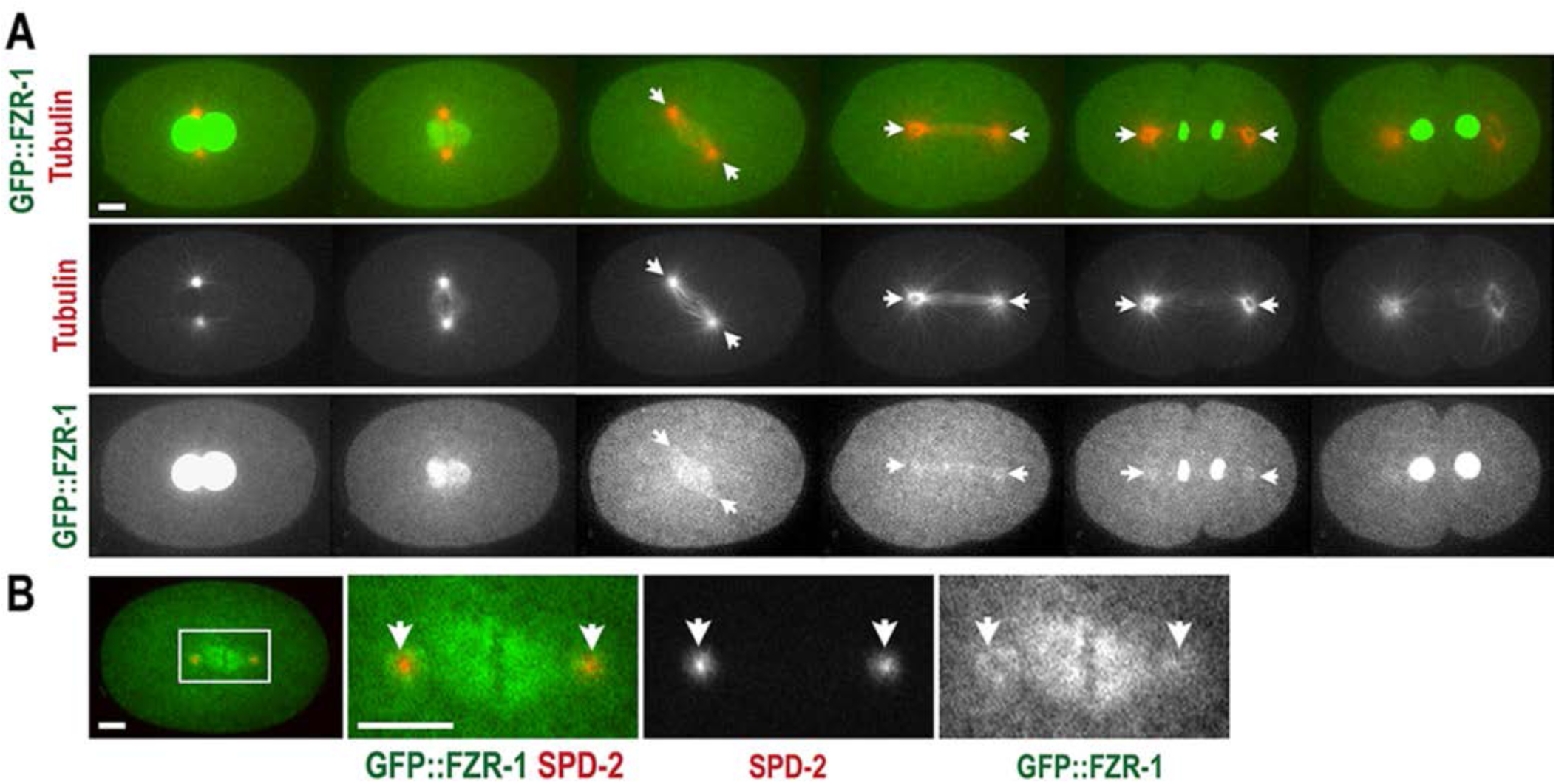
Subcellular localization of FZR-1 during the first cell cycle. (A) Still images from time-lapse movie of an embryo expressing GFP∷FZR-1 and mCherry∷tubulin. Movie was acquired at 1 min interval. GFP∷FZR-1 localizes at nuclei, mitotic spindles and centrosomes (arrows). Expression of mCherry∷tubulin used as a subcellular land-marker. (B) Embryo expressing GFP∷FZR-1 and mCherry∷SPD-2 displays that GFP∷FZR-1 localizes to mitotic spindles and centrosomes (arrows) that co-localize with mCherry-SPD-2, a centrosome marker. Bar, 5 μm.

### FZR-1 might function as a part of the APC/C complex to regulate centrosome duplication

Given that FZR-1 is a conserved co-activator of the APC/C, an E3 ubiquitin ligase, we hypothesized that FZR-1 functions as a part of the APC/C complex in centrosome assembly. If so, depleting other APC/C subunits should have a similar effect that loss of FZR-1 had on the *zyg-1(it25)* mutant. To examine how other core subunits of the APC/C complex might affect *zyg-1(it25)* mutants, we mated the *zyg-1(it25)* strain with *mat-3(or180)* mutants for the core APC8/CDC23 subunit (Golden *et al.* 2000), and *emb-1(hc57)* mutants for the conserved subunit APC16 in the *C. elegans* APC/C complex (Kops *et al.* 2010; Green *et al.* 2011; Shakes *et al.* 2011). By generating double homozygote mutants, we assayed for bipolar spindle formation and embryonic viability in *zyg-1(it25); mat-3(or180)* and *zyg-1(it25); emb-1(hc57)* double homozygous mutants (Figure 3, Table 1). At the restrictive temperature 24°, *zyg-1(it25); mat-3(or180)* double mutant embryos exhibit a 9-fold increase in bipolar spindle formation (81.8±14.3%), compared to *zyg-1(it25)* single mutant embryos (9.1±8.8%) during the second mitosis (Figure 3A). Consistently, 5% of *zyg-1(it25); mat-3(or180)* double mutants produce viable progeny while 100% of *zyg-1(it25)* or *mat-3(or180)* single mutant progeny die at 24° (Table 1). In support of our results, the *mat-3(bs29)* allele has been reported as a genetic suppressor of *zyg-1* (Miller *et al.* 2016). Furthermore, we observed that the *emb-1* mutation suppresses the centrosome duplication phenotype of *zyg-1(it25)* mutants at the semi-restrictive temperature 22.5°. While 45.5±11.9% of *zyg-1(it25)* embryos form bipolar spindles, 79.1±12.4% of *zyg-1(it25); emb-1(hc57)* double mutant embryos produce bipolar spindles (Figure 3A). We, however, observed no significant restoration of embryonic viability in *zyg-1(it25); emb-1(hc57)* double mutants (*p*=0.691) compared to *zyg-1(it25)* single mutants (Table 1), presumably due to the strong embryonic lethality by the *emb-1(hc57)* mutation itself (Kops *et al.* 2010; Shakes *et al.* 2011). Our results indicate that the APC/C complex functions to suppress the phenotype of *zyg-1(it25)* mutants. Therefore, FZR-1 might function as a component of the APC/C complex to regulate centrosome duplication in early *C. elegans* embryos.

**Figure 3.**
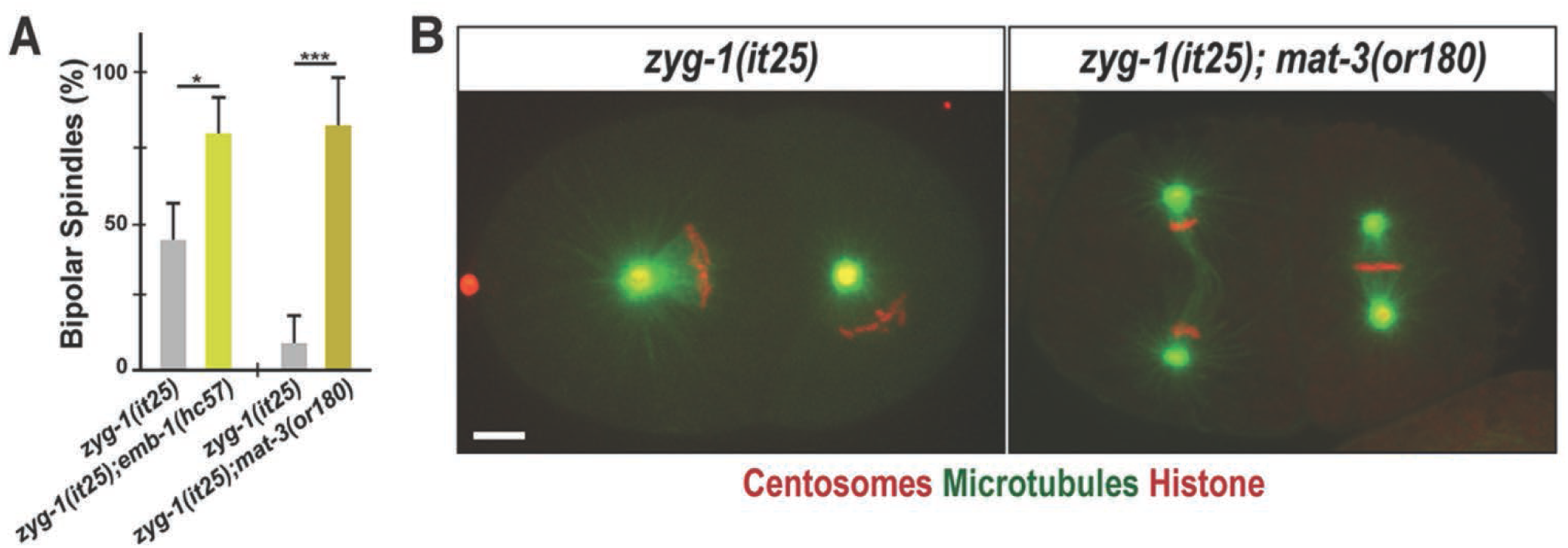
Inactivating the APC/C restores bipolar spindle formation to *zyg-1(it25)*. (A) Quantification of bipolar spindle formation during the second cell cycle. At 22.5°, there is an increase in bipolar spindle formation in *zyg-1(it25); emb-1(hc57)* double mutants (79.1±12.4%, n=228, *p*=0.03), compared to *zyg-1(it25)* single mutants (45.5±11.9%, n=238). At 24°, *zyg-1(it25); mat-3(or180)* double mutants assembled bipolar spindles at a significantly higher percentage (81.8±14.3%, n=78, *p*<0.001) than *zyg-1(it25)* embryos (9.1±8.8%, n=144). n is the number of blastomeres. \**p*<0.05, \*\*\**p*<0.001 (two-tailed t-test). (B) Still images of embryos expressing GFP∷β tubulin, mCherry∷γ-tubulin (centrosome marker) and mCherry∷histone raised at 24° illustrate monopolar spindle formation in the *zyg-1(it25)* embryo, and bipolar spindle formation in the *zyg-1(it25); mat-3(or180)* double mutant embryo. Bar, 5 μm.

### Loss of FZR-1 results in elevated SAS-5 levels

Next we wanted to understand how FZR-1 contributes to centrosome duplication. Since FZR-1 appears to function through the APC/C complex in centrosome assembly, we hypothesized that the APC/C^FZR-1^ specifically targets one or more centrosome regulators for ubiquitin-mediated degradation. If that is the case, depleting FZR-1 should protect substrates from degradation leading to accumulation of target proteins. To identify a direct substrate of APC/C^FZR-1^ that regulates centrosome assembly, we utilized the conserved FZR-1 co-activator specific recognition motif, KEN-box, to screen for a potential substrate (Glotzer *et al.* 1991; Pfleger and Kirschner 2000; Song and Rape 2011). The KEN-box appears to be the major degron motif that APC/C^FZR-1^ recognizes in centrosome duplication (Strnad *et al.* 2007; Tang *et al.* 2009; Arquint and Nigg 2014; Meghini *et al.* 2016). In human cells, HsSAS-6, STIL/SAS-5, and CPAP/SAS-4 contain KEN-box motif, and APC/C^Cdh1/FZR-1^ targets these proteins for ubiquitin-mediated proteolysis, thereby preventing extra centrosomes (Strnad *et al.* 2007; Tang *et al.* 2009; Arquint *et al.* 2012; Arquint and Nigg 2014). The *Drosophila* APC/C^Fzr/Cdh1/FZR-1^ is also shown to target Spd2 for destruction through direct interaction with a KEN-box (Meghini *et al.* 2016). Interestingly in *C. elegans*, a putative KEN-box motif is present in SAS-5 but none in SAS-4 and SAS-6, which indicates an evolutionary divergence between humans and nematodes.

Protein stabilization by the *fzr-1* mutation might lead to increased levels of a centrosome-associated substrate, which may compensate for impaired ZYG-1 function at the centrosome. In *C. elegans*, SAS-5 is the only core centrosome duplication factor containing a KEN-box, which suggests SAS-5 as a potential target of the APC/C^FZR-1^. If the APC/C^FZR-1^ targets SAS-5 directly through KEN-box for ubiquitin-mediated proteolysis, inhibiting FZR-1 should protect SAS-5 from degradation leading to SAS-5 accumulation. To examine how the *fzr-1* mutation affected SAS-5 stability, we immunostained embryos with anti-SAS-5, and quantified the fluorescence intensity of centrosome-associated SAS-5 (Figure 4A and 4B). As ZYG-1 is required for SAS-5 localization to centrosomes, hyper-accumulation of SAS-5 might compensate for partial loss-of-function of ZYG-1, thereby restoring centrosome duplication to *zyg-1(it25)* mutants. In fact, our quantitative immunofluorescence revealed that *fzr-1(bs31)* embryos exhibit a significant increase (1.41±0.42 fold; *p*<0.001) in centrosomal SAS-5 levels at the first anaphase, compared to wild-type (Figure 4B). Consistently, compared to *zyg-1(it25)* single mutants, *zyg-1(it25); fzr-1(bs31)* double mutant embryos exhibit a 1.48-fold increase (*p*<0.001) in centrosome-associated SAS-5 levels (Figure 4B). Indeed, centrosomal SAS-5 are restored to near wild-type levels in *zyg-1(it25); fzr-1(bs31)* double mutants (0.95±0.44 fold; *p*=0.003). We, however, observed no significant changes in centrosomal TBG-1 (γ-tubulin) levels in *fzr-1(bs31)* mutants (Figure S1).

**Figure 4.**
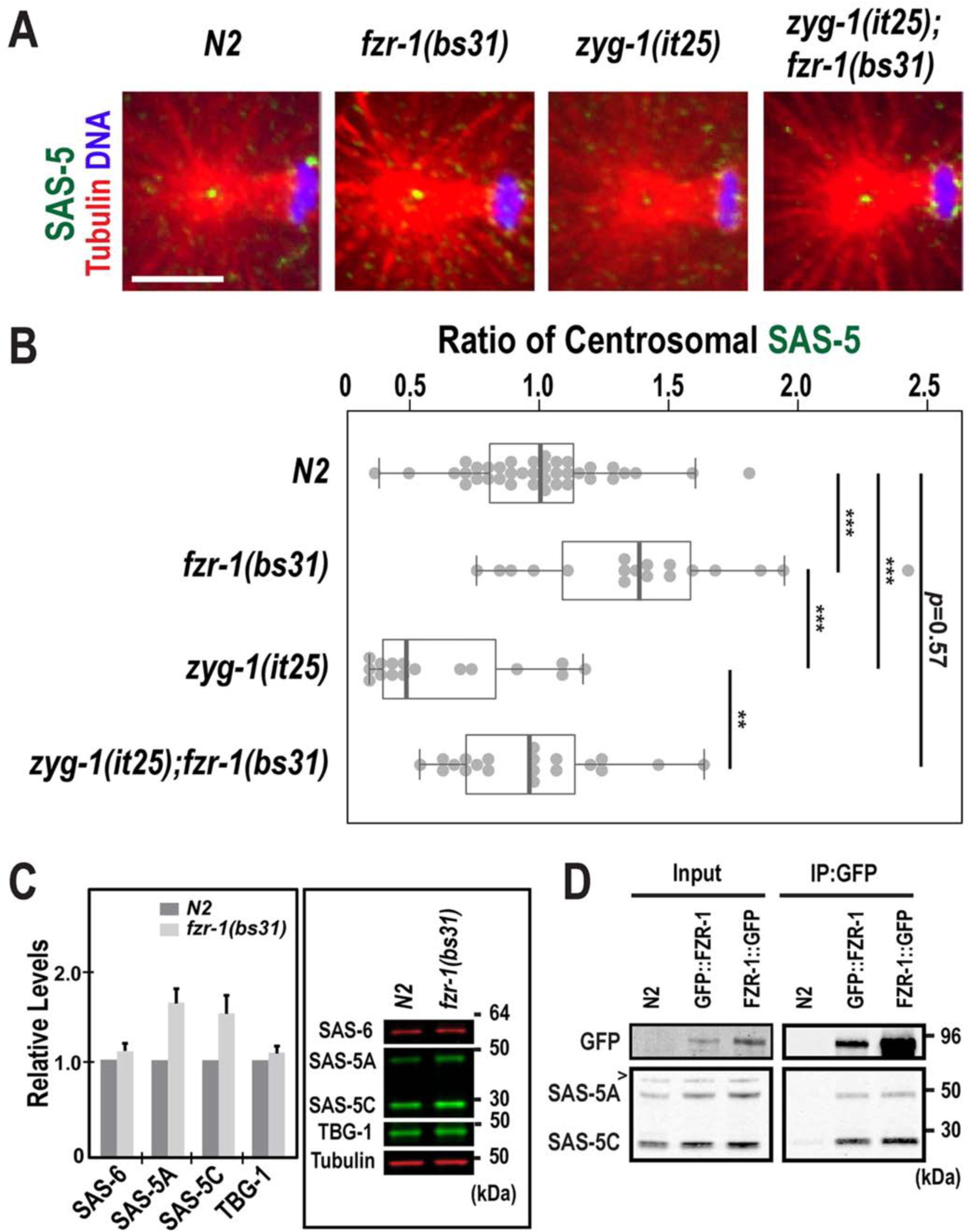
Loss of FZR-1 results in elevated SAS-5 levels. (A) Images of centrosomes stained for SAS-5 (green) at the first anaphase. Bar, 5 μm. (B) Quantification of centrosome-associated SAS-5 levels at the first anaphase. SAS-5 levels are normalized to the average fluorescence intensity in wild-type centrosomes. *fzr-1(bs31)* embryos exhibit increased levels of centrosomal SAS-5 (1.41±0.42 fold, n=18; *p*<0.001) relative to wild-type embryos (1.00±0.28 fold, n=38). In *zyg-1(it25); fzr-1(bs31)* double mutants, centrosomal SAS-5 levels are restored to near wild-type levels (0.95±0.44 fold, n=20; *p*=0.003), compared to *zyg-1(it25)* embryos that show decreased levels of centrosomal SAS-5 (0.64±0.28 fold, n=16). n is the number of centrosomes. Each dot represents a centrosome. Box ranges from the first through third quartile of the data. Thick bar indicates the median. Dashed line extends 1.5 times the inter-quartile range or to the minimum and maximum data point. \*\**p*<0.01, \*\*\**p*<0.001 (two-tailed t-test). (C) Quantitative western blot analyses show that (left panel) *fzr-1(bs31)* mutant embryos possess increased levels of both SAS-5 isoforms, SAS-5A (1.56±0.16 fold) and SAS-5C (1.48±0.19 fold), compared to wild-type (*N2*) embryos. However, there were no significant differences in levels of either SAS-6 (1.09±0.08 fold) or TBG-1 (1.08±0.07 fold) between *fzr-1(bs31)* mutant and wild-type embryos. Four biological samples and eight technical replicates were used. Average values are presented and error bars are SD. (right panel) Representative western blot using embryonic lysates from *fzr-1(bs31)* mutants and *N2* animals. Tubulin was used as a loading control. (D) Immunoprecipitation (IP) using anti-GFP suggests that FZR-1 physically interacts with SAS-5. Both SAS-5 isoforms (SAS-5A, SAS-5C) co-precipitate with GFP∷FZR-1 or FZR-1∷GFP. Wild-type (*N2*) embryos were used as a negative control of IP. ~1% of total embryonic lysates was loaded in the input lanes. ‘>’ indicates a non-specific detection by the SAS-5 antibody.

Elevated protein levels might influence centrosome-associated SAS-5 levels in *fzr-1(bs31)* mutants. To determine how inhibition of the APC/C^FZR-1^ affected overall protein levels, we performed quantitative western blot analysis using embryonic protein lysates and antibodies against centrosome proteins (Figure 4C). Our data indicate that *fzr-1(bs31)* embryos possess increased SAS-5 levels (~1.5-fold), relative to wild-type embryos, while the levels of SAS-6 and TBG-1 are not significantly affected in *fzr-1(bs31)* mutants (Figure 4C). Our observation on the SAS-6 levels in *fzr-1(bs31)* mutants is consistent with previous work by Miller *et al.*, 2016, showing no increase in SAS-6 levels by the *mat-3(bs29)*/APC8 mutation that inhibits the APC/C function. These results suggest that *C. elegans* utilizes a different mechanism to control SAS-6 levels, unlike Human SAS-6 that is regulated by the APC/C-mediated proteolysis (Strnad *et al.* 2007). Furthermore, our immunoprecipitation suggests a physical interaction between SAS-5 and FZR-1 in *C. elegans* embryos (Figure 4D), supporting that SAS-5 might be a direct substrate of the APC/C^FZR-1^. Consistent with our results in this study, prior study has shown that inhibiting the 26S proteasome leads to increased levels of SAS-5 (Song *et al.* 2011). Thus, SAS-5 levels are likely to be controlled through the ubiquitin-proteasome system.

Collectively, our data show that the *fzr-1* mutation leads to a significant increase in both cellular and centrosomal levels of SAS-5, suggesting that the APC/C^FZR-1^ might control SAS-5 levels via ubiquitin-mediated proteasomal degradation to regulate centrosome assembly in the *C. elegans* embryo.

### Mutation of the KEN-box stabilizes SAS-5

If the APC/C^FZR-1^ directly targets substrates for destruction via the conserved KEN-box, mutating this motif should cause substantial resistance to the ubiquitination-mediated degradation. To determine whether the APC/C^FZR-1^ targets SAS-5 through the KEN-box motif, we mutated the KEN-box at the endogenous *sas-5* locus. By using CRISPR/CAS-9 mediated genome editing (Paix *et al.* 2015), we generated mutant lines (*sas-5^KEN-to-^*^3A^) carrying alanine substitutions of the SAS-5 KEN-box (Figure 5A). The *sas-5^KEN-to-3A^* mutant embryo exhibits no obvious cell cycle defects or embryonic lethality (Table 1), consistent with *fzr-1* mutants (Kemp et al., 2007). *sas-5^KEN-to-3A^* animals exhibit a slightly reduced (~80%) and irregular distribution of brood size within the population (Figure S2). Reduced brood size and slow growth phenotypes were previously reported in *fzr-1* mutant alleles (Fay *et al.* 2002; Kemp *et al.* 2007).

**Figure 5.**
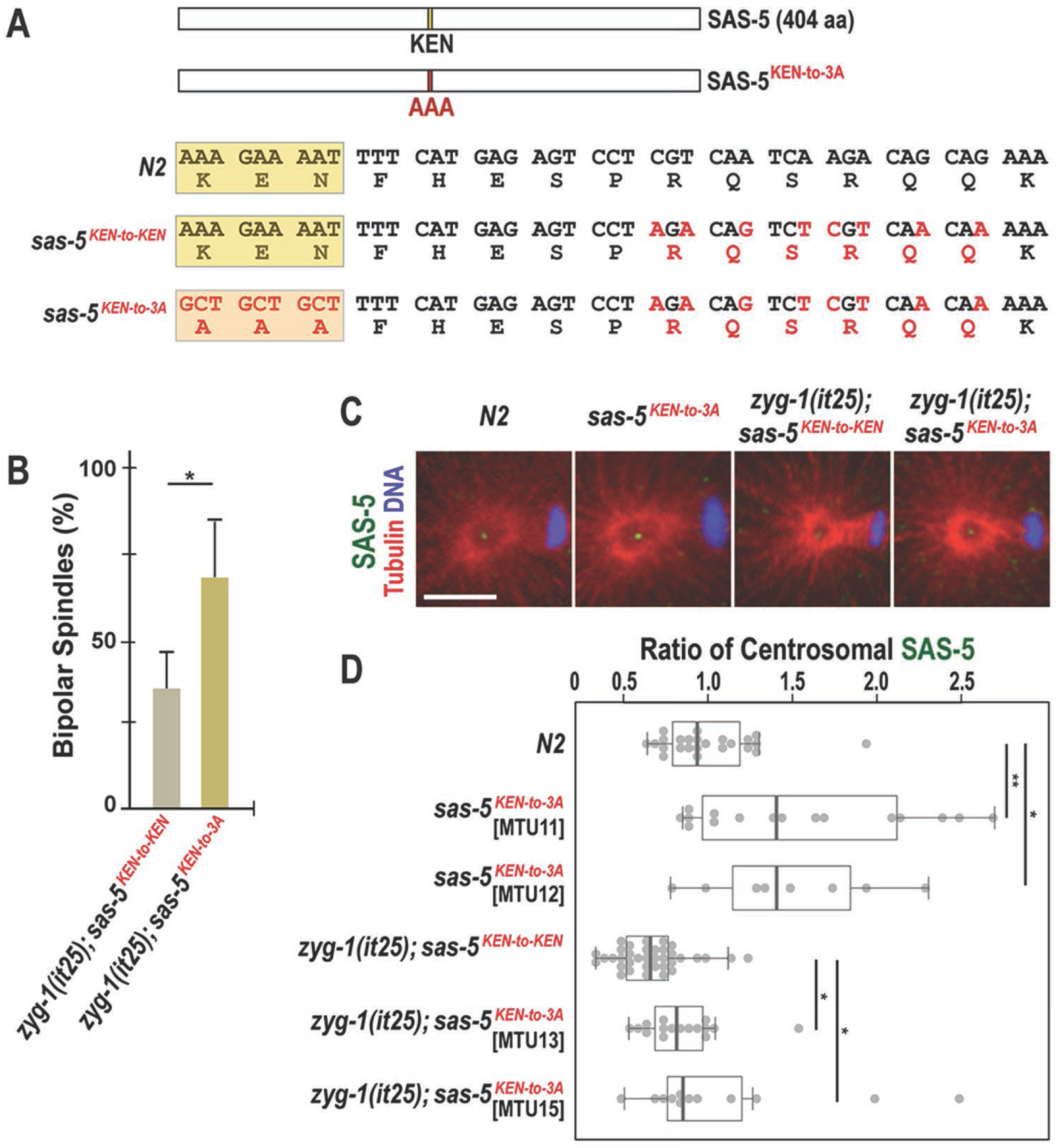
Mutation of the SAS-5 KEN-box leads to increased SAS-5 levels at centrosomes and restores centrosome duplication to *zyg-1(it25)* mutants. (A) SAS-5 contains a KEN-box (aa 213-216) motif. Mutations (red) are introduced at multiple sites to make alanine substitutions (AAA; 3A) for the KEN-box and additional silent mutations for the CRISPR genome editing (see methods and materials). The KEN-box is highlighted in yellow. Note that the *sas-5*^*KEN-to-KEN*^ mutation contains the wild-type SAS-5 protein. (B) Quantification of bipolar spindle formation during the second cell cycle in *zyg-1(it25); sas-5*^*KEN-to-KEN*^ and *zyg-1(it25); sas-5*^*KEN-to-3A*^ embryos at 22.5°. *zyg-1(it25); sas-5*^*KEN-to-3A*^ double mutant embryos produce bipolar spindles at a higher rate (67.5±16.3%, n=124, *p*=0.02) than *zyg-1(it25); sas-5*^*KEN-to-KEN*^ controls (35.1±10.7%, n=164). n is the number of blastomeres. Average values are presented and error bars are SD. (C) Centrosomes stained for SAS-5 (green) during the first anaphase. Bar, 5 μm. (D) Quantification of centrosomal SAS-5 levels during the first anaphase. We used two independently generated *sas-5*^*KEN-to-3A*^ mutant lines to quantify SAS-5 levels (MTU11 and 12, Table S1). SAS-5 levels at centrosomes are normalized to the average fluorescence intensity in wild-type centrosmes. Mutating the SAS-5 KEN-box leads to increased levels of centrosomal SAS-5 in both MTU11 (1.54±0.63 fold, n=16; *p*=0.04) and MTU12 (1.48±0.50 fold, n=8; *p*=0.03), compared to wild type (1.00±0.29 fold; n=24 centrosomes). Consistently, there are a significant increase in centrosomal SAS-5 levels in both *zyg-1(it25); sas-5*^*KEN-to-3A*^ double mutant lines (MTU13: 0.85±0.24 fold, n=16; *p*=0.01 and MTU15: 1.09±0.59 fold; n=12; *p*=0.03), compared to *zyg-1(it25); sas-5*^*KEN-to-KEN*^ control that contains reduced levels of centrosomal SAS-5 (0.67±0.20 fold; n=36 centrosomes). n is the number of centrosomes. Each dot represents a centrosome. Box ranges from the first through third quartile of the data. Thick bar indicates the median. Dashed line extends 1.5 times the interquartile range or to the minimum and maximum data point. \**p*<0.05, \*\**p*<0.01 (two-tailed t-test).

Next, we asked how the *sas-5^KEN-to-3A^* mutation affected *zyg-1(it25)* mutants. If the APC/C^FZR-1^-mediated proteolysis of SAS-5 accounts for the suppression of *zyg-1, sas-5^KEN-to-3A^* mutants should mimic the *fzr-1* mutation that suppresses *zyg-1* mutants. By mating the *sas-5^KEN-to-3A^* mutant with *zyg-1(it25)* animals, we tested whether the *sas-5^KEN-to-3A^* mutation could genetically suppress *zyg-1* mutants, by assaying for embryonic viability and centrosome duplication (Table 1, Figure 5B). For the *zyg-1(it25)* mutant control in this experiment, we used the strain MTU14 [*zyg-1(it25); sas-5^KEN-to-KEN^*, Table S1] that contains the equivalent modifications, except the KEN-box, to the *sas-5^KEN-to-3A^* mutation (Figure 5A, see methods and materials). At the semi-restrictive temperature 22.5°, *zyg-1(it25)*; *sas-5^KEN-to-3A^* animals lead to a 7.7-fold increase in the frequency of viable progeny (35.3±9.2%; *p*<0.0001), compared to *zyg-1(it25); sas-5^KEN-to-KEN^* mutant controls (4.6±4.0%) (Table 1). Consistently, *zyg-1(it25)*; *sas-5^KEN-to-3A^* embryos exhibit successful bipolar spindle assembly at a significantly higher rate (67.5±16.3%; *p*=0.02) than *zyg-1(it25); sas-5^KEN-to-KEN^* embryos (35.1±10.7%) at the two-cell stage (Figure 5B). These results suggest that the *sas-5^KEN-to-3A^* mutation does partially restore embryonic viability and centrosome duplication to *zyg-1(it25)* mutants at 22.5°. However, at the restrictive temperature 24° where the *fzr-1* mutation shows a strong suppression (Table1, Figure 1B), both *zyg-1(it25)*; *sas-5^KEN-to-3A^* double mutants and *zyg-1(it25); sas-5^KEN-to-KEN^* mutant animals result in 100% embryonic lethality (Table 1). *zyg-1(it25)*; *sas-5^KEN-to-3A^* embryos (14.7% bipolar, n=68) grown at 24° show only minor effect on centrosome duplication compared to *zyg-1(it25); sas-5^KEN-to-KEN^* control embryos (7.6% bipolar, n=66). The data obtained at 24° reveal that the *sas-5^KEN-to-3A^* mutation results in much weaker suppression to *zyg-1(it25)* mutants than the *fzr-1* mutation, suggesting that the SAS-5 KEN-box mutation does not generate the equivalent impact that results from the *fzr-1* mutation. If SAS-5 is the only APC/C^FZR-1^ substrate that contributes to the suppression of *zyg-1* mutants, the *fzr-1* or KEN-box mutation might influence SAS-5 stability differently. In this scenario, FZR-1 might target SAS-5 through KEN-box and additional recognition motifs (e.g., D-box), causing a greater effect on SAS-5 stability than the KEN-box mutation alone. To examine how the KEN-box mutation affected SAS-5 stability, we measured the fluorescence intensity of SAS-5 at centrosomes by quantitative immunofluorescence (Figure 5C and 5D). At 22.5° where the *sas-5^KEN-to-3A^* mutation restores centrosome duplication and embryonic viability to *zyg-1(it25)*, *sas-5^KEN-to-3A^* mutants exhibit a significant increase in centrosome-associated SAS-5 levels (~1.5-fold, *p*<0.001), compared to wild-type (Figure 5C and D). Consistently, *zyg-1(it25)*; *sas-5^KEN-to-3A^* embryos display ~1.4-fold (*p*=0.002) increased SAS-5 levels at centrosomes, compared to *zyg-1(it25); sas-5^KEN-to-KEN^* control embryos that contain reduced centrosomal SAS-5 levels (Figure 5D). Notably, *zyg-1(it25)*; *sas-5^KEN-to-3A^* embryos exhibit centrosomal SAS-5 levels nearly equivalent (~0.97 fold) to those of wild-type embryos (Figure 5D). As a control, we also quantified centrosomal TBG-1 levels but saw no changes between *sas-5^KEN-to-3A^* mutants and the wild-type (Figure S1). Furthermore, we examined overall SAS-5 levels by quantitative western blot, finding that relative to wild-type embryos, *sas-5^KEN-to-3A^* mutant embryos possess ~1.5-fold increased SAS-5 levels (Figure S3). Together, our quantification data reveal that the *sas-5*^*KEN-to-3A*^ or *fzr-*1 mutation leads to nearly equivalent fold change (~1.5-fold) in both cellular and centrosome-associated SAS-5 levels (Figure 4B, 4C, 5D and S3). Together, these results suggest that APC/C^FZR-1^ directly targets SAS-5 in a KEN-box dependent manner to control SAS-5 turnover, and that SAS-5 stabilization by blocking proteolysis results in elevated SAS-5 levels at the centrosome, partially contributing to the suppression of the *zyg-1(it25)* mutation. In human cells, APC/C^Cdh1^ recognizes a KEN-box to regulate the levels of STIL, the homolog of *C. elegans* SAS-5, and STIL depleted of the KEN-box leads to accumulation of STIL protein, and centrosome amplification (Arquint and Nigg 2014). While we do not observe extra centrosomes by the SAS-5 KEN-box mutation, our data show that that APC/C^FZR-1^ controls SAS-5 stability via the direct recognition of the conserved degron motif, KEN-box, to regulate centrosome duplication in *C. elegans* embryos, suggesting a conserved mechanism for regulating SAS-5 levels between humans and nematodes.

Interestingly, although either inhibiting FZR-1 or mutating KEN-box influences SAS-5 stability at a comparable level, we observe a notable difference in the suppression level by these two mutations. Weaker suppression by the *sas-*5^*KEN-to-3A*^ mutation suggests that the APC/C^FZR-1^ might target additional substrates that cooperatively support the *zyg-*1 suppression. In this scenario, APC/C^FZR-1^ might target other centrosome proteins outside core duplication factors through the conserved degron motifs, such as destruction (D)-box and KEN-box (Glotzer *et al.* 1991; Pfleger and Kirschner 2000). Alternatively, APC/C^FZR-1^ might target additional core centrosome factors through other recognition motifs other than KEN-box, such as D-box (Glotzer *et al.* 1991) or unknown motif in the *C. elegans* system. In humans and flies, APC/C^Cdh1/Fzr^ has been shown to regulate the levels of STIL/SAS-5, Spd2, HsSAS-6 and CPAP/SAS-4 (Strnad *et al.* 2007; Tang *et al.* 2009; Arquint and Nigg 2014; Meghini *et al.* 2016). While *C. elegans* homologs of these factors, except SAS-5, lack a KEN-box, all five centrosome proteins contain at least one putative D-box. An intriguing possibility, given the strong genetic interaction observed between *fzr-1* and *zyg-1*, is that ZYG-1 could be a novel substrate of APC/C^FZR-1^. Additional work will be required to understand the complete mechanism of APC/C^FZR-1^- dependent regulation of centrosome duplication in *C. elegans*. In summary, our study shows the APC/C^FZR-1^-dependent proteolysis of SAS-5 partially contributes to the suppression of the *zyg-*1 mutants, and we report that FZR-1 functions as a negative regulator of centrosome duplication in *C. elegans*.

## Acknowledgements

We thank members of Song lab (Naomi Haque, Brittany Rettig and Michael Stubenvoll) for their technical support, Kevin O’Connell and Andy Golden for RNAi and worm stains. We especially thank WormBase and the *Caenorhabditis* Genetics Center (CGC). WormBase is supported by grant U41 HG002223 from the National Human Genome Research Institute at the US National Institutes of Health, the UK Medical Research Council and the UK Biotechnology and Biological Sciences Research Council. The CGC (St. Paul, MN), is funded by the National Institutes of Health Office of Research Infrastructure Programs (P40 OD010440).

## Author Contributions

J.C.M. and M.H.S. designed the experiments and wrote the manuscript. J.C.M. and M.H.S. performed quantifications of confocal imaging and protein levels from western blots. J.C.M., L.E.D. M.M.K., and M.H.S. performed experiments and provided data.

---

**Figure S1.**
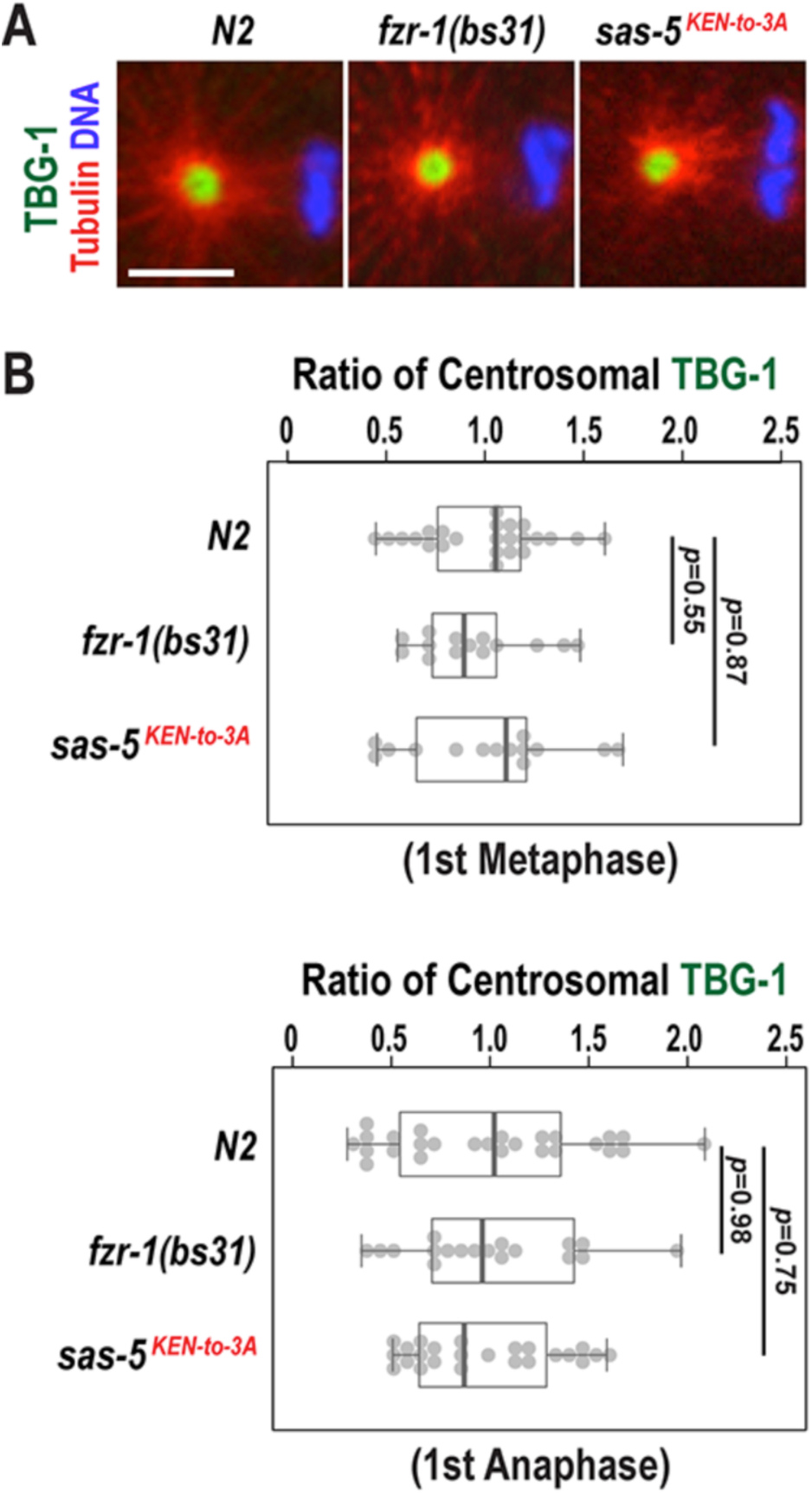
Centrosome-associated TBG-1 levels are unaffected in *fzr-1(bs31)* and *sas-5^KEN-to-3A^* mutant embryos. (A) Centrosomes stained for TBG-1 (green) at the first metaphase. Bar, 5μm. (B) Quantification of TBG-1 levels at centrosomes during the first mitosis. TBG-1 levels are normalized to the average fluorescence intensity in wild-type (*N2*) embryos. At the first metaphase, *fzr-1(bs31)* (0.94±0.28 fold, n=14; *p*=0.55) and *sas-5*^*KEN-to-3A*^ mutants (1.02±0.40 fold, n=14; *p*=0.87) have comparable centrosomal TBG-1 levels to wild-type (1.00±0.30 fold, n=24). At the first anaphase, centrosome-associated TBG-1 levels in both *fzr-1(bs31)* (0.99±0.42 fold, n=18; *p*=0.98) and *sas-5*^*KEN-to-3A*^ (0.96±0.36 fold, n=24; *p*=0.75) mutant embryos are similar to those of wild-type (1.00±0.52 fold, n=26). n is the number of centrosomes. Each dot represents a centrosome. Boxes ranges from the first through third quartile of the data. Thick bar indicates the median. Dashed line extends 1.5 times the inter-quartile range or to the minimum and maximum data point.

**Figure S2.**
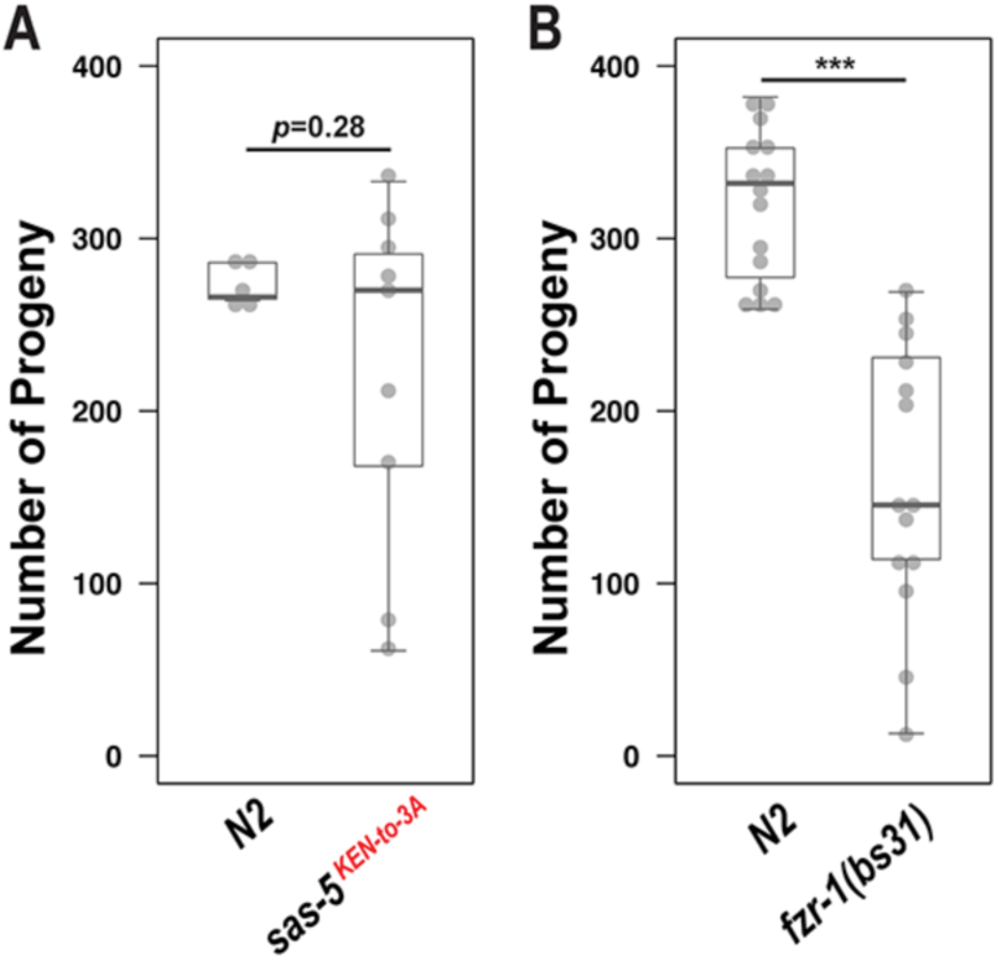
Brood size in *fzr-1(bs31)* and *sas-5*^*KEN-to-3A*^ mutants. (A) *fzr-1(bs31)* mutants produce reduced brood size (158.4±78.4, n=14 hermaphrodites; *p*<0.001) compared to wild-type animals (319.6±43.9, n=15 hermaphrodites) grown at 24°. Note that *fzr-1(bs31)* mutants produce a wide range of distribution in brood size among 14 animals tested, which is also seen in *sas-5^KEN-to-3A^* mutants. (B) *sas-5^KEN-to-3A^* mutants display a slight reduction in brood size (222.7±99.7, n=9 hermaphrodites; *p*=0.28) compared to wild-type controls (273.4±11.5, n=5 hermaphrodites) grown at 24°. Compared to wild-type animals, *sas-5*^*KEN-to-3A*^ mutant animals produce highly irregular number of progeny in the population of nine animals tested under the same condition. Each dot represents the total number of progeny produced by a single animal. Box ranges from the first through third quartile of the data. Thick bar indicates the median. Dashed line extends 1.5 times the inter-quartile range or to the minimum and maximum data point.

**Figure S3.**
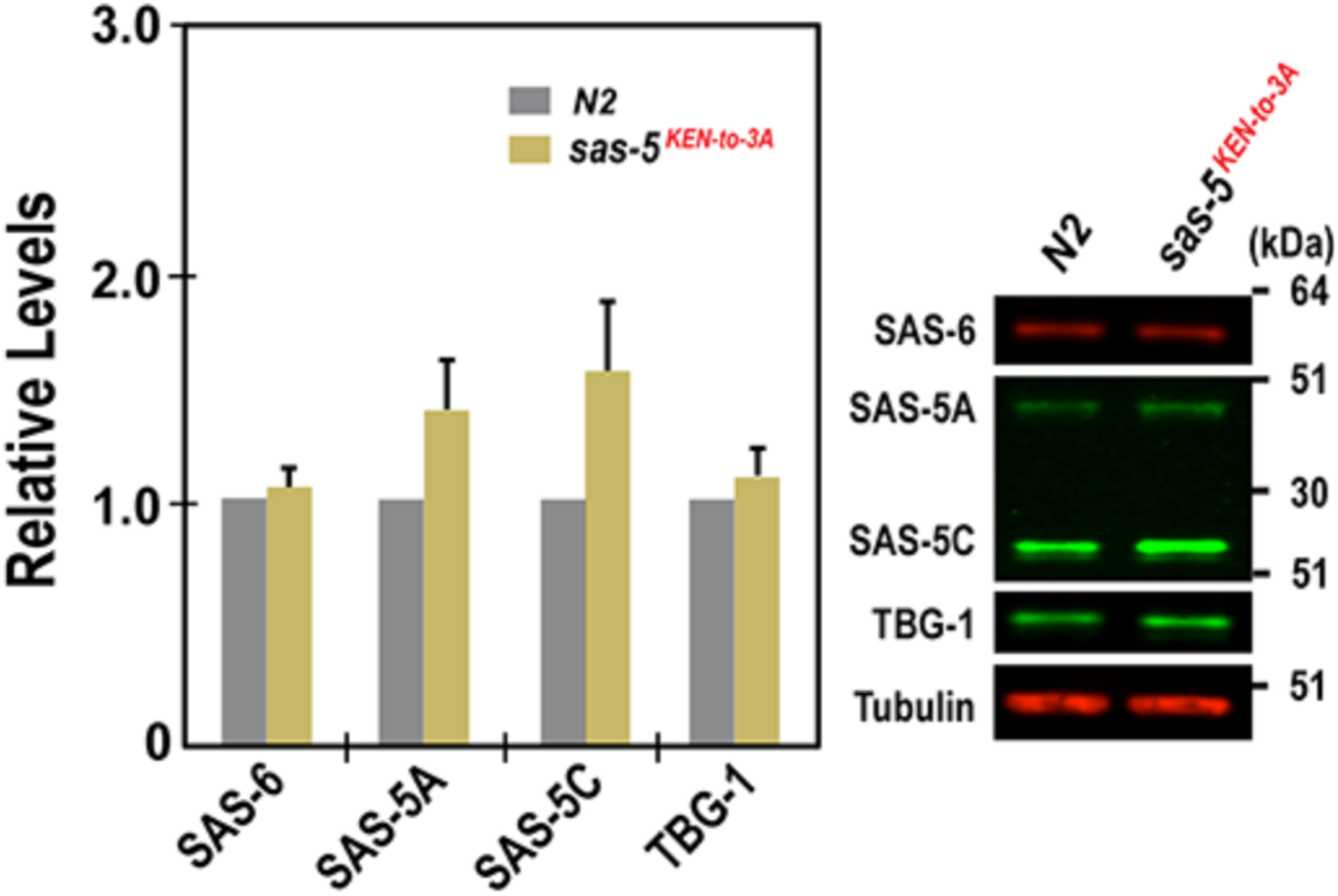
SAS-5 levels are increased in *sas-5*^*KEN-to-3A*^ mutants. Quantitative western blot reveal that (left panel) ***sas-5^KEN-to-3A^*** mutant embryos contain increased levels of both SAS-5 isoforms, SAS-5A (1.36±0.20 fold) and SAS-5C (1.52±0.28 fold), compared to wild-type (*N2*) embryos. In contrast, there were no significant changes in either SAS-6 (1.05±0.04 fold) or TBG-1 (1.09±0.11 fold) levels between *sas-5^KEN-to-3A^* mutant and wild-type embryos. Four biological samples and six technical replicates were used for the statistical analysis. Average values are presented and error bars are SD. (right panel) Representative western blot using embryonic lysates from *sas-5*^*KEN-to-3A*^ mutants and *N2* animals. Tubulin was used as a loading control.

**Table S1.** List of *C. elegans* Strains Used in This Study

| Name | Genotype | Origin |
| --- | --- | --- |
| N2 | *wild-type* | CGC |
| CB4856 | *wild-type, Hawaiian variant* | CGC |
| CB120 | *unc-4(e120) II* | Brenner 1974 |
| CB128 | *dpy-10(e128) II* | Brenner 1974 |
| MJ57 | *emb-1(hc57) III* | Schierenberg *et al.* 1980 |
| MTU6 | *fzr-1(bs31) II* | This study |
| MTU7 | *mat-3(or180) III* | Golden *et al.* 2000 |
| MTU8 | *zyg-1(it25) II; mat-3(or180) III* | This study |
| MTU9 | *zyg-1(it25) II; emb-1(hc57) III* | This study |
| MTU10 | *unc-119(ed3) III; [unc-119(+); fzr-1p::fzr-1::gfp::fzr-1 3'UTR]* | This study<br>Sarov *et al.* 2012 |
| MTU11 | *sas-5(mhs357) [sas-5<sup>KEN-to-3A</sup>] V* | This study |
| MTU12 | *sas-5(mhs358) [sas-5<sup>KEN-to-3A</sup>] V* | This study |
| MTU13 | *zyg-1(it25) II; sas-5(mhs359) [sas-5<sup>KEN-to-3A</sup>] V* | This study |
| MTU14 | *zyg-1(it25) II; sas-5(mhs361) [SAS-5<sup>KEN-to-KEN</sup>] V* | This study |
| MTU15 | *zyg-1(it25) II; sas-5(mhs362) [sas-5<sup>KEN-to-3A</sup>] V* | This study |
| OC13 | *zyg-1(or409) II* | Kemp *et al.* 2007 |
| OC14 | *zyg-1(it25) II* | Kemphues *et al.* 1988 |
| OC130 | *zyg-1(it25); fzr-1(bs38) II* | Kemp *et al.* 2007 |
| OC190 | *unc-119(ed3) III; bsIs6 [unc-119(+), pie-1p::gfp::fzr-1]* | This study |
| OC201 | *zyg-1(it25); fzr-1(bs31) II* | Kemp *et al.* 2007 |
| OC481 | *unc-119(ed3) III; bsIs15[pNP99; unc-119(+), tbb-1p::mCherry::tbb-2::tbb-2 3'UTR]* | Gift from O'Connell Lab<br>Medley *et al.* 2017 |
| OC740 | *bsSi15[pKO109; unc-119(+), spd-2p::spd-2::mCherry::spd-2 3'UTR] I* | Peel *et al.* 2017 |
| SA250 | *tjIs54[pie-1p::GFP::tbb-2 + pie-1p::2xmCherry::tbg-1 + unc-119(+)]; tjIs57[pie-1p::mCherry::his-48 + unc-119(+)]* | Toya *et al.* 2010 |

**Table S2.**
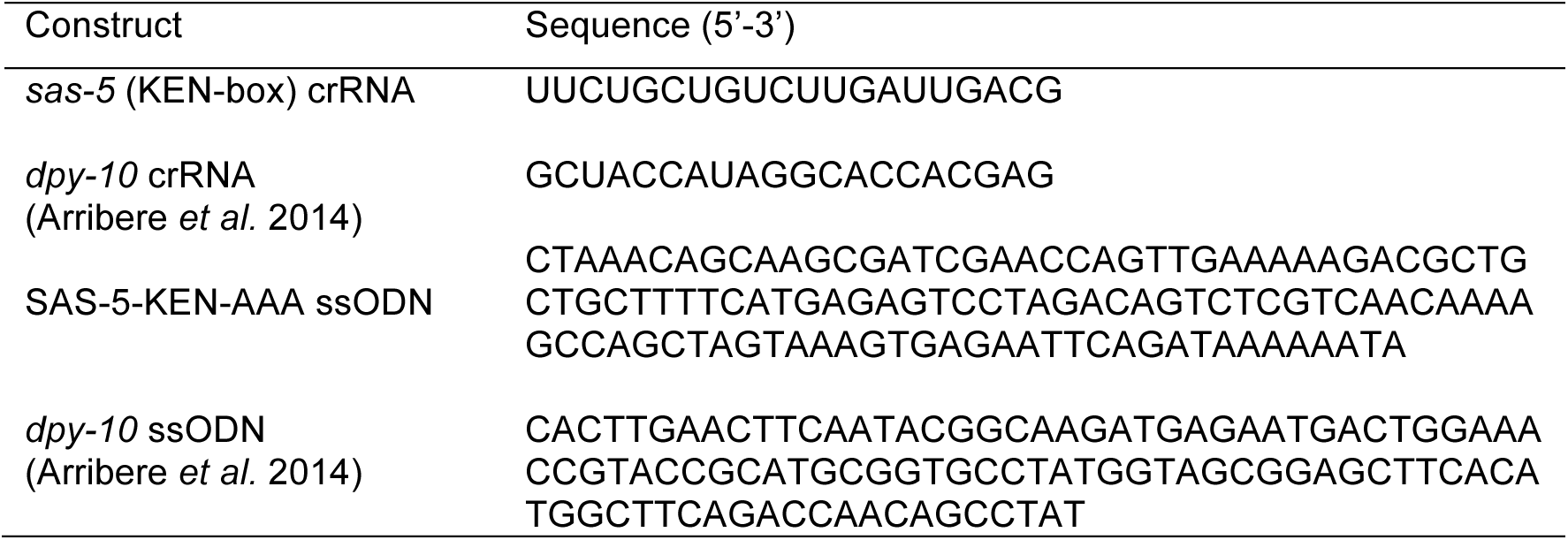
List of Oligonucleotides for CRISPR/Cas9 Genome Editing

